# A stromatoporoid-like sediment-agglutinating sponge from microbialites of Cambrian Stage 4

**DOI:** 10.1101/2025.09.25.678569

**Authors:** Cui Luo, Joachim Reitner, Zhixin Sun, Kai Chen

## Abstract

Many extant demosponges, especially the non-spicular taxa, incorporate foreign detritus in their spongin or chitinous skeletons. This supposed energy-efficient way of skeleton construction is, however, rarely known in the fossil record. In this study, based on careful petrographic analyses using standard optical microscopy, cathodoluminescence and fluorescence microscopy, and a three-dimensional reconstruction, a fossil from the Cambrian Stage 4 of Xuzhou City, Jiangsu Province, China, is established as a sediment-agglutinating sponge *Psammolectospongia beiwangensis* gen. et sp. nov. These fossils encrust stromatolitic microbialites and are often surrounded by *Girvanella* mats. This succession of microbialites and sponge fossils was developed in a turbulent hydrodynamic condition after the deposition of edgewise conglomerates. *P. beiwangensis* possesses labyrinthine skeletons with an architecture similar to that of some stromatoporoids, such as *Stromatopora* and *Syringostromella*, although dissepiments are absent in the Cambrian fossil. The skeleton of *P. beiwangensis* is mainly composed of packed silt-to sand-sized detritus grains and skeletal fragments of other animals, while in some places, it can contain various proportions of micrites. The morphology and skeletal composition of *P. beiwangensis* indicate an affinity to an agglutinating demosponge, which mainly constructed its skeleton using agglutinated and incorporated sediments, while the spongin skeleton was able to be biomineralized like that of *Vaceletia*. This study is the first report of a sponge fossil that mainly built its skeleton using foreign detritus, and it expands our knowledge about the physiology and ecological behaviors of early Paleozoic demosponges.

## 1. Introduction

The fossil record of early sponges, especially those during the Neorpterozoic-early Palaeozoic transitional interval, is critical for reconstructing the early evolutionary trajectory of animals (e.g., Carlisle et al., 2024; Sperling et al., 2010). Despite molecular fossils (Love et al., 2020, 2009), fossil sponges are mainly recognized based on the skeletal morphology that is preserved in the rock record and comparable with that of known poriferan taxa. The most well-known sponge skeleton fossils include scattered sponge spicules (e.g., Bengtson, 1986), articulated spicular sponge fossils (e.g., Rigby, 1986), and the massive carbonate basal skeleton of coralline/hypercalcified sponges (Hartman and Goreau, 1970). The organic fibrous skeleton of non-spicular sponges may also have the chance to be fossilized in shale and carbonates (e.g., Luo et al., 2022; Rigby, 1986; Wu et al., 2024), although the involved taphonomic mechanisms require experimental explanation, and in each specific example, it requires scrutiny to confirm the non-spicular-sponge origin and exclude other provenances which produce similar structures (Luo, 2023; Luo et al., 2022; Neuweiler et al., 2023).

Additional to these biologically secreted skeletons, sponges can also agglutinate and incorporate foreign detritus to reinforce their skeletal frameworks. This behavior is not limited to, but especially common among the extant non-spicular demosponges (Wiedenmayer 1989, Cook, 2007; Schönberg, 2016). Sponges adopting this strategy have managed to flourish in environments with high sedimentation rates and can rapidly build skeletons, possibly with a lower energy cost relative to biomineralization (Schönberg, 2016). This strategy seems to be especially useful for early fossil sponges, especially when they needed to survive and develop in an ancient oxygen-depleted ocean. Seilacher (1994) proposed the concept of “sand sponges” when discussing the possible presence of sand skeletons in Ediacaran-Cambrian metazoans. However, active sediment agglutination and incorporation are so far rarely confirmed in fossil sponges, except for the Jurassic example of the *Calciagglutispongia yabei* Reitner, 1992. In this study, we describe a stromatoporoid-like fossil from Cambrian Stage 4 that exhibits the habit of sediment incorporation.

## 2. Materials and methods

The investigated specimens were collected from the Beiwang section in the northeast of Beiwang village (34°14′4.85″N, 117°6′22.3″E), Xuzhou City, Jiangsu Province, North China. Xuzhou is located at the southeastern margin of the North China Block (Fig. 1A). The subdivision and nomenclature of local Cambrian successions vary among schools of geologists (Fang, 1997). Following the lithostratigraphic frame presented in Zhu et al. (2021), they are composed of Houjiasha, Mantou, Zhangxia, Chaomidian, and Sanshanzi formations in ascending order (Fig. 1B).

**Fig. 1.**
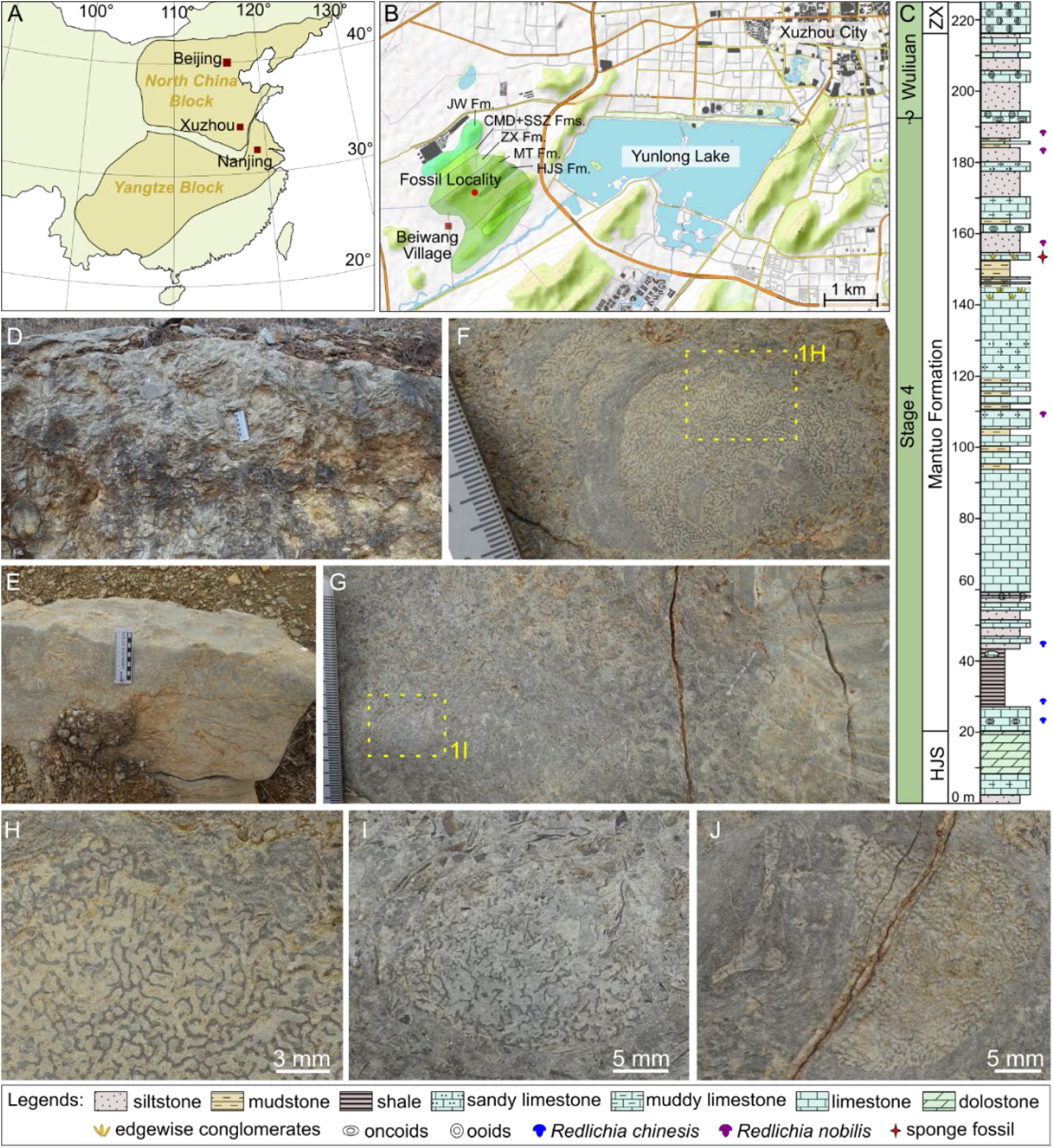
Location, stratigraphy, and field observation of the fossil locality. (A) Location of Xuzhou in China, modified after Zhu et al. (2021). (B) The fossil locality in Xuzhou City. The distribution of the Cambrian strata is modified after the Regional Geological Survey Team of Jiangsu Geological Bureau (1977). (C) The stratigraphic log of the Mantou Formation near the Beiwang Village, drawn after the description in Chen (2020). (D) Edgewise conglomerates at the outcrop, plan view. (E) Side view of the boulder containing labyrinthine fossils shown in F–J. The fossil horizon is on top of the edgewise conglomerates. (F–J) Labrynnthine fossils and their surrounding host fabrics. Areas in the yellow dashed rectangles are enlarged in the noted figures, respectively. Abbreviations in B and C: CMD, Chaomidian Formation; HJS, Houjiashan Formation; JW, Jiawang Formation; MT, Mantou Formation; SSZ, Sanshanzi Formation; ZX, Zhangxia Formation.

The Mantou (‘Manto’) Formation is a well-known Cambrian lithostratigraphic unit in North China and is composed of purple mudstone at its type section (Zhang, 1996). In north Jiangsu Province, the Mantou Formation is characterized by purple shale intercalated with marl, limestone, dolostone, and sandstone (Fang, 1997). In the study section, an echinoderm species has been reported from the purple shale in the lower Mantou Formation (Wang et al., 2024). The stromatoporoid-like fossils were recovered from the upper part of the Mantou Formation, within the *Redlichia nobilis* Zone (Chen, 2020)(Fig. 1C), which was correlated to the middle part of Stage 4, Cambrian Series 2 (Yoon et al., 2022; Zhu et al., 2019).

The fossil locality is on the margin of an expanding graveyard, and the fossils were collected from boulders that had fallen only a few meters away from their original position. The correlation between the boulders and the bedrock can be confirmed, as the fossil-bearing layer develops immediately on top of edgewise conglomerates (Fig. 1E), which are comparable with the *in situ* edgewise conglomerate bed (Fig. 1D) in components, conglomerate sizes, color, and texture. The fossils are exposed on an erosional surface (Fig. 1E–J), but unfortunately, this surface is not preserved in the ambient *in situ* exposures.

Two fossil-containing blocks were sampled and processed into 24 thin sections and a few polished slabs, containing 6 investigated fossil individuals (hereafter numbered as SPF-1 to SPF-6). The thin sections were initially studied using a Nikon SMZ18 microscope coupled with a Nikon DSRi2 camera, then investigated using a Zeiss Axio Imager 2 fluorescence microscopy and a CITL Mk5-2 cold cathodoluminescence (CL) system coupled to a petrographic microscope. In order to visualize the 3-D structure of the fossil, part of SPF-1 was serially ground using an EXAKT 400 CS grinder. An Epson V370 scanner was used to collect images of the serial planes. One hundred planes were prepared with an average inter-plane distance of 31 ± 3 μm (Supplementary Material 1A). The obtained images were processed using Adobe Photoshop CS6 for picture alignment and ImageJ 1.54p for segmentation (Supplementary Material 1B and C). In order to improve the visualization, the segmented image stack was interpolated by 100 planes using the scaling tool in ImageJ 1.54p (Supplementary Material 1D). Finally, the re-scaled image stack was rendered using Voreen 5.2.0 (voreen.uni-muenster.de) (Supplementary Material 1E and F). All analyses except for CL were conducted at the Nanjing Institute of Geology and Palaeontology (NIGPAS). CL was conducted at Nanjing University. All specimens are deposited at NIGPAS.

## 3. Results

The studied fossils appear as labyrinthine patches growing on top of stromatolitic microbialites, which in turn, encrust a thin bed of grainstone composed of rounded, sand-sized clasts (Fig. 2A and C). The staining using alizarin red S showed that the labyrinthine fossils and the microbialites are all preserved as calcite. The components in this succession will be described one by one below. Before formally establishing the sponge affinity in section 4.1, the stromatoporoid-like fossils will be called “labyrinthine fossils”, and the supposed skeletons called “partitions”.

**Fig. 2.**
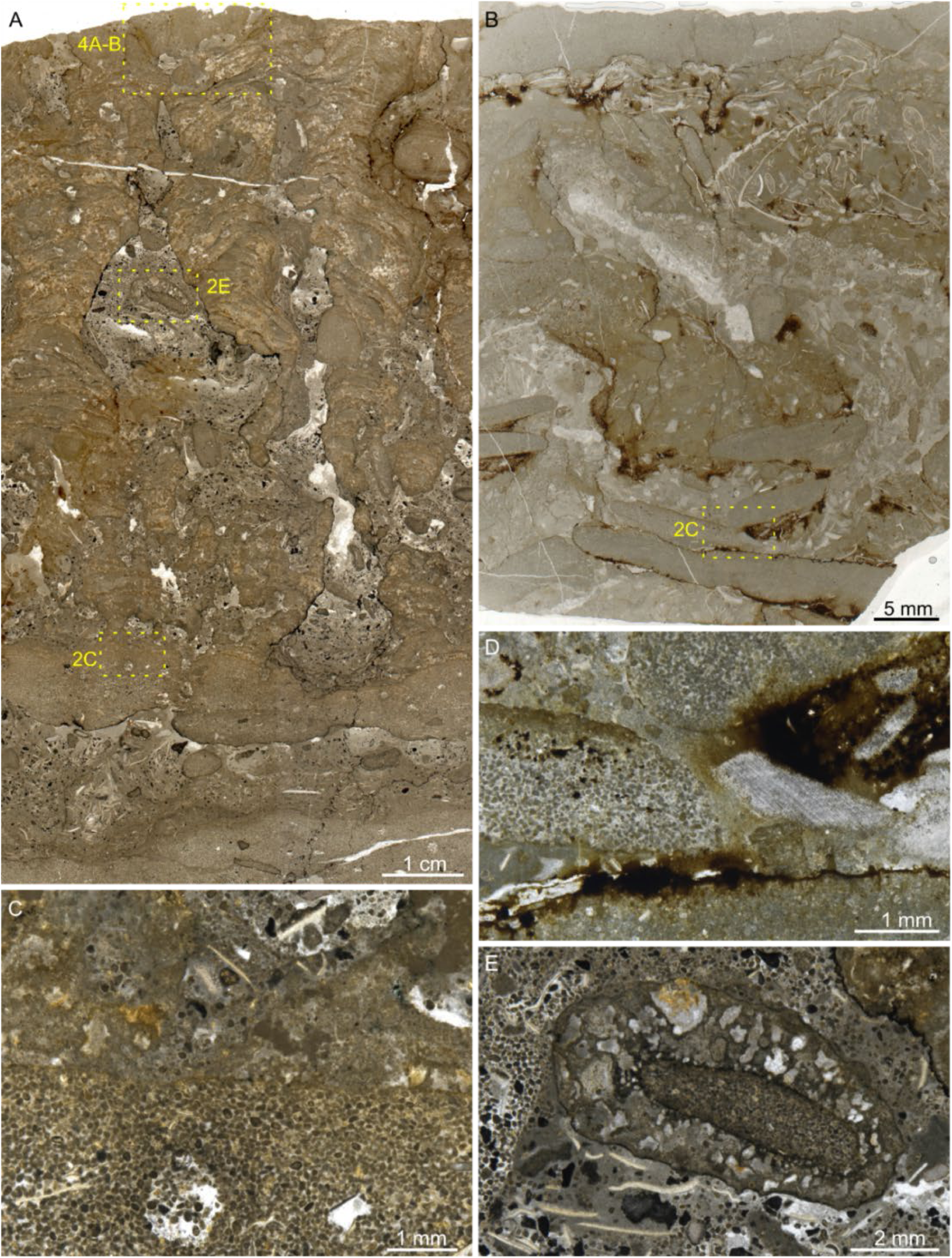
The depositional successions underlying the labyrinthine fossil SPF-3. (A) The vertical cross-section of the labrynthine fossil SPF-3, the underlying stromatolitic microbialite, and the two grainstone beds. (B) Vertical cross-section of the edgewise conglomerates underlying the succession shown in Fig. 2A. (C) Close-up of an area in Fig. 2A, showing the contact between the stromatolite and the grainstone bed. (D) A close view of the flat pebbles in Fig. 2B. (E) A peculiar oncolite composed of a core of a flat pebble enveloped in a clotted fabric. Thin section numbers: xz-15 for A, C, E; xz-06 for B, D.

### 3.1. The substrate of the microbialite

The grainstone bed immediately underlying the stromatolitic microbialites is a typical hardground (Demicco and Hardie, 1995; Flügel, 2010), as it shows a distinct and smooth boundary against the encrusting microbialites and exhibits syndepositional erosions in places (Figs. 2A and 2C). However, syndepositional burrows, which are partly filled by the same detrital grains as those in the host rock, are scattered in this bed (Fig. 2A and C), indicating the activity of animals in the short interval before the lithification of the grainstone.

A second grainstone layer lies below the described one and is separated from the latter by a 1–2 cm thick floatstone (Fig. 2A). Further below this second grainstone bed is the edgewise conglomerate (Fig. 2B and D). This grainstone layer, thicker and horizontally more stable than the first one, overlies the edgewise conglomerate pebbles (Fig. 2B). The flat pebbles show a similar composition to the described grainstone beds, although some pebbles contain a higher proportion of mud or larger grains (Fig. 2D).

However, despite the succession observed in thin sections and slabs, the field observation has shown that the edgewise conglomerate extends up to the weathered bedding surface where the labyrinthine fossils are exposed (Fig. 1E–G). Therefore, it seems that the surface of the edgewise conglomerate bed was originally undulating, and the microbialites and labyrinthine fossils were developed or selectively preserved only in the depressions.

### 3.2. The stromatolitic microbialites

In the analyzed samples, the microbialites underlying the labyrinthine fossils are columns up to 9 cm tall and mostly 1–2 cm wide. The columns show an anastomosed branching habit and are separated by intercolumn spaces filled with skeletal and rock fragments of different sizes, as well as sparitic cements (Fig. 2A, C, E, 3A). Notably, among the intercolumn fillings, there is a peculiar sort of oncolite which often exceeds 2 mm in size and consists of a core of a grainstone pebble and a thick envelope with mottled, probably microbial fabrics (Fig. 2E).

**Fig. 3.**
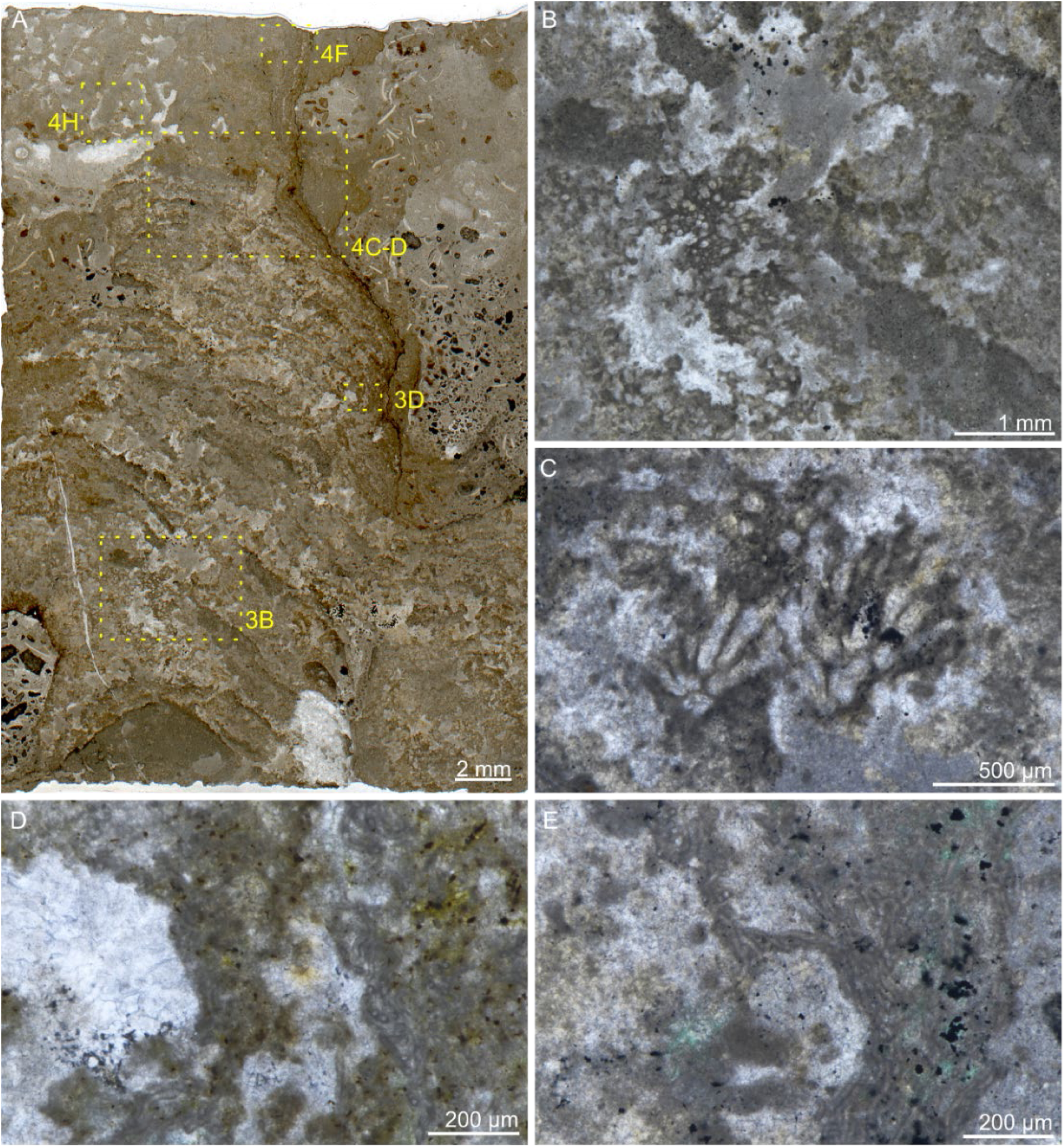
Characteristics of the stromatolitic microbialites underlying the labyrinthine fossil SPF-2. (A) A vertical thin section showing SPF-2 encrusing over the stromatolitic microbialite. (B and C) *Hedstroemia*-type calcimicrobes that destroy laminations in the microbialites. (D and E) Primary cavities formed by the growth of *Girvanella* in the microbialites. Thin section numbers: xz-05a-1 for A, B, D; xz-05a-2 for C; xz-05b-1 for E.

The microbialites show crude laminations in a general view, but each lamination is discontinuous and dominantly consists of clotted elements (Fig. 2A, 3A). This mesostructure is, indeed, intermediate between a typical stromatolite and a typical thrombolite (Aitken, 1967; Grey and Awramik, 2020; Kennard and James, 1986; Riding, 2000). However, since the laminated pattern is most prominent at the mesostructure scale (i.e., centimetric scale) (Shapiro, 2000), these microbialites are preferentially called stromatolites in this study.

The crude lamination reflects the stepwise accretion of the microbialites, while the clotted fabric seems to be conferred by multiple processes. First, although microbe fossils are rarely preserved in these microbialites, in some places, the bundles of entangled *Girvanella* filaments are observed forming primary cavities (Fig. 3D and E). This growth mode was known from other Neoproterozoic and Phanerozoic analogs, when microbially produced gas bubbles were trapped in mats mainly composed of filamentous microbes (Lee and Riding, 2016; Mata et al., 2012). Second, the growth of some *Hedstroemia*-type calcimicrobes (Liu et al., 2017; Riding, 1991) interrupts the laminae (Fig. 3B and C). Third, the development of laminae can also be disrupted by increased debris and syndepositional erosion (upper-left corner of the stromatolite in Fig. 3A).

### 3.3. The labyrinthine fossils

In the plan view on a weathered rock surface, these fossils appear as irregular patches with labyrinthine partitions, often surrounded by a gray wall (Fig. 1F, H–J). In vertical cross-section, they are found encrusting on stromatolitic microbialites (Fig. 2A, 3A, 4A–D, 5C and F). The geometry of the partitions can be as isotropically labyrinthine as in the plan view (Fig. 3A and 4A) or be more elongated along the top-base axis (Fig. 5C). In all studied specimens, the partitions are more readily recognizable in reflected light than in transmitted light (Fig. 4A–D).

**Fig. 4.**
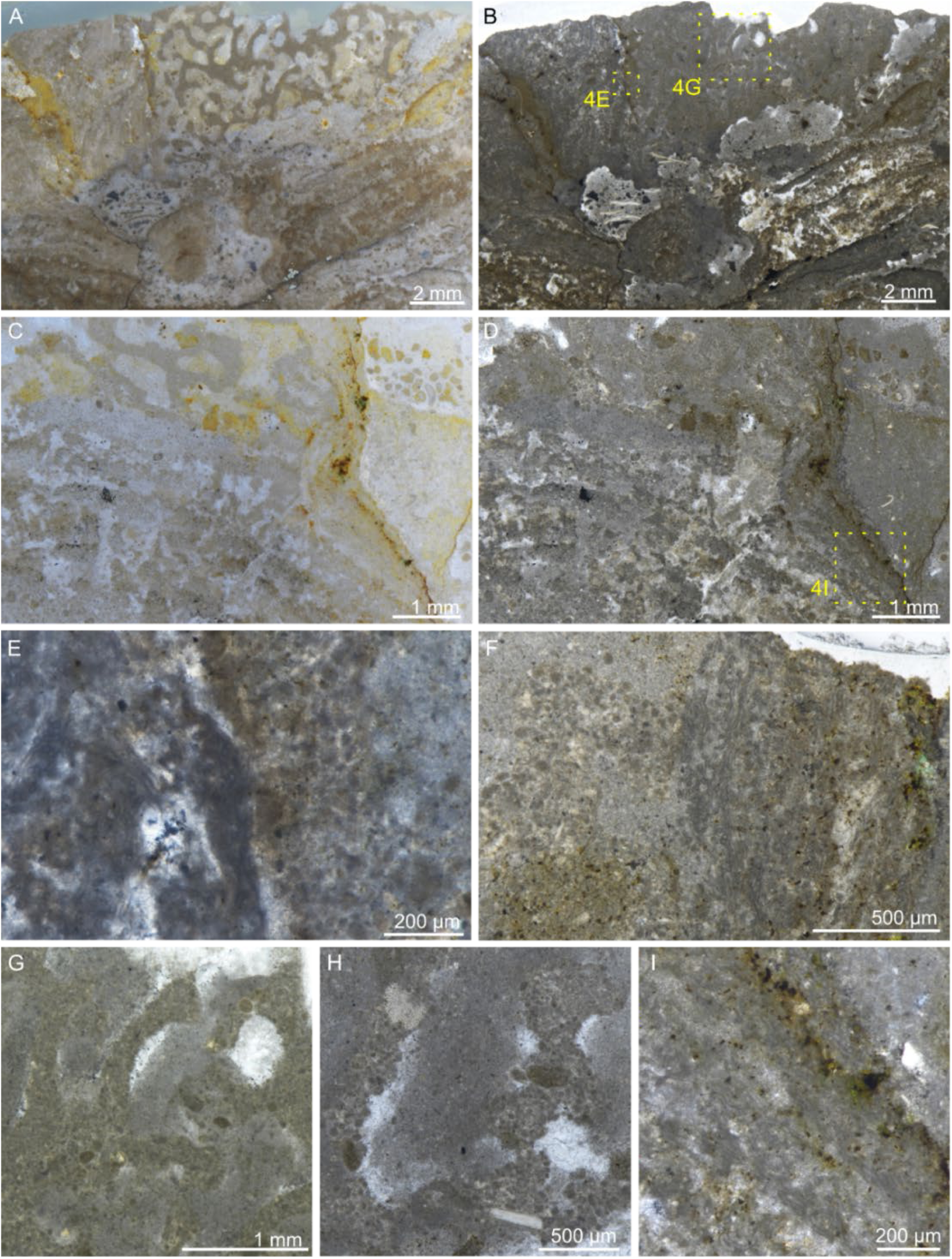
Architecture and composition of the partitions of the labyrinthine fossils. (A and B) Vertical cross-section of SPF-3 in reflected (A) and transmitted light (B), respectively. (C and D) Vertical cross-section of SPF-2 in reflected (C) and transmitted light (D), respectively. (E and F) The distinct contact between the partitions and the *Girvanella* mats surrounding the labyrinthine fossils. (G and H) The labyrinthine partitions composed of packed sand-to silt-sized allochthonous grains. (I) The calcified *Girvanella* mat in the outermost layer of the stromatolite below SPF-2. Thin section numbers: xz-15 for A, B, E, G; xz-05a-1 for C, D, F, H, I.

**Fig. 5.**
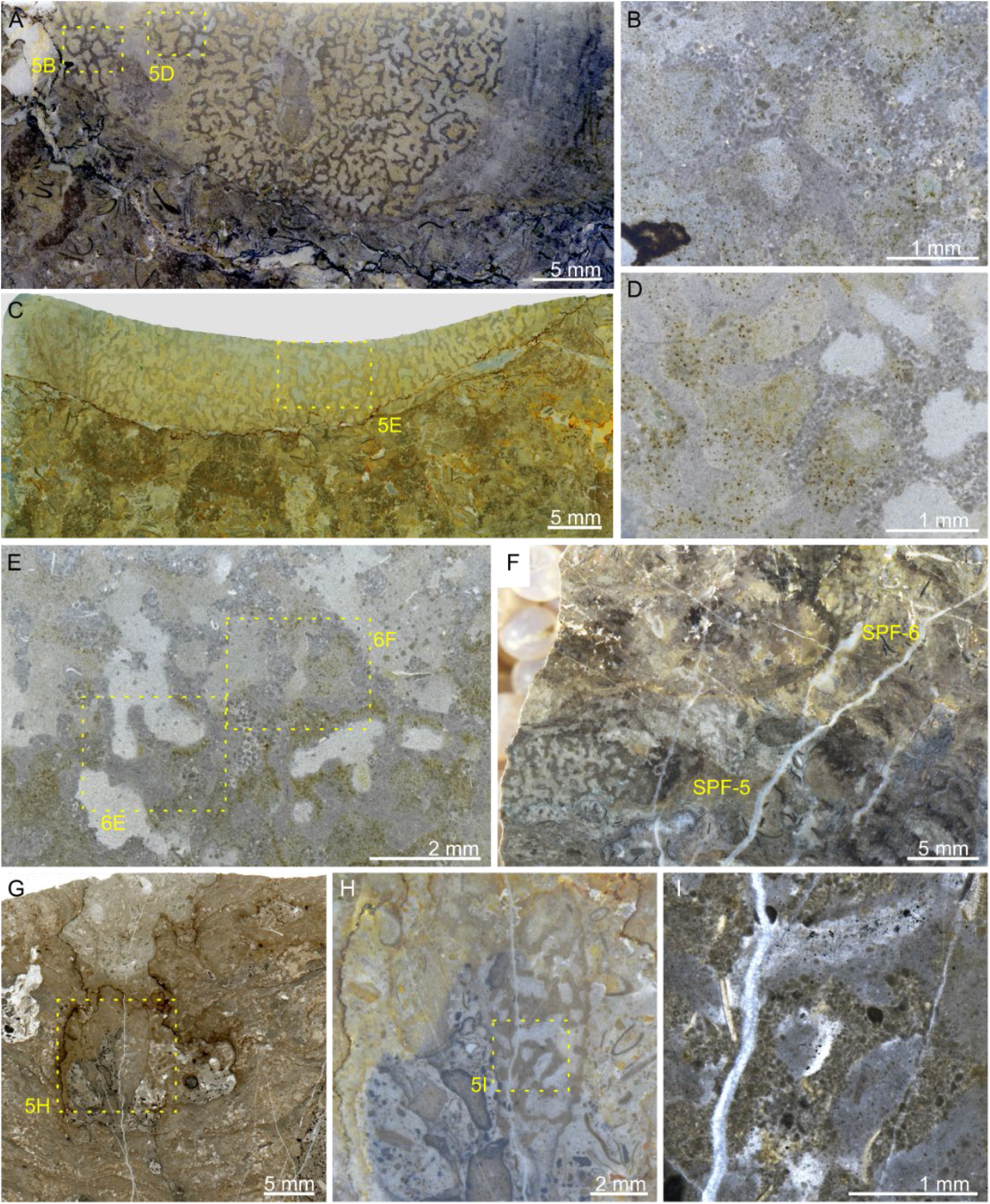
Morphology and composition of SPF-1, SPF-4, SPF-5, and SPF-6. (A) Thin section scan showing the plan view of SPF-1. (B, D, E) Parts of the thin sections in A and C, showing the transition between micritic and detrital compositions in the partitions. (C) Thin section scan showing the vertical view of SPF-1. (F) The slab showing SPF-5 and SPF-6 intercalated in the microbialite. (G and H) The thin section of SPF-4 in transmitted (G) and reflected light (H). (I) The detrital composition of SPF-4. Thin section and slab numbers: xz-03 for A, B, D; xz-02 for C, E; xz-15-slab for G; xz-14 for G–I.

The contact between the labyrinthine fossil and the stromatolite is often eroded or altered by pressure solution (Fig. 2A, 3A, 4A and B, 5C and F). However, in some places, the contact is almost intact and shows a distinct immediate contact between them (Fig. 4C and D). The wall surrounding the labyrinthine fossils observed in the field (Fig. 1F, H, J) turns out to be a mat woven by *Girvanella* (Fig. 4E and F). This *Girvanella* mat can be continuous with the outermost layers of the underlying stromatolite (Fig. 4C, D, I), while it always shows a distinct boundary against the labyrinthine partitions, never intruding into the latter (Fig. 4E and F).

The partitions are around 100–230 μm thick, dominantly composed of cemented silt-to sand-sized detritus grains and animal skeletal fragments (Fig. 4E–H). The interspaces measured between subparallel partitions are mostly around 300–600 μm wide, filled by micrites, sparitic cements, and grains similar to those in the partitions. Notably, the density of detritus grains distributed in the partition interspaces is much lower than that accumulated outside of the fossil, and the average size of particles is also obviously smaller (Fig. 3A, 4A–D, 5A). The partitions, although composed of packed grains, can show very smooth and distinct margins (Fig. 4G). Moreover, in the specimen SPF-1, part of the partitions is composed of micrites, which grade into areas composed of packed grains (Fig. 5B and D). Interestingly, in the vertical cross-section, the proportion of micritic components seems to be higher in the lower part of the specimen (Fig. 5C and E).

Despite these specimens observable from the top of the microbialites, SPF-4, SPF-5, and SPF-6 were found in polished slabs, in the middle of the microbialites. SPF-4 is preserved in an intercomlumn cavity of the thrombolitic stromatolites (Fig. 5G–I), possibly representing a reworked example. SPF-5 and SPF-6 encrust on and are intercalated in the microbialites (Fig. 5F).

The cathodoluminescence behavior of the studied materials seems to be primarily associated with crystal sizes: the micritic parts are generally brighter and the sparitic parts darker, regardless of the origins of the carbonates (Fig. 6A–D). In the labyrinthine partitions, the difference between the dull cements and the luminescent grains is noticeable (Fig. 6B and D). For fluorescence microscopy, previous studies suggested that blue incident light produces the best result for carbonate petrology (Dravis and Yurewicz, 1985), and this is consistent with our observation. Therefore, in Fig. 6E–J, only the results from the FITC mode are illustrated. The results show that the partitions are generally non-fluorescent, although some inter-particle cements can be as bright as the fillings in the inter-partition spaces (Fig. 6E–G). This method performs well to differentiate the labyrinthine partitions from the interspace fillings, even when they look similar in a standard optical microscope (Fig. 6E– G).

**Fig. 6.**
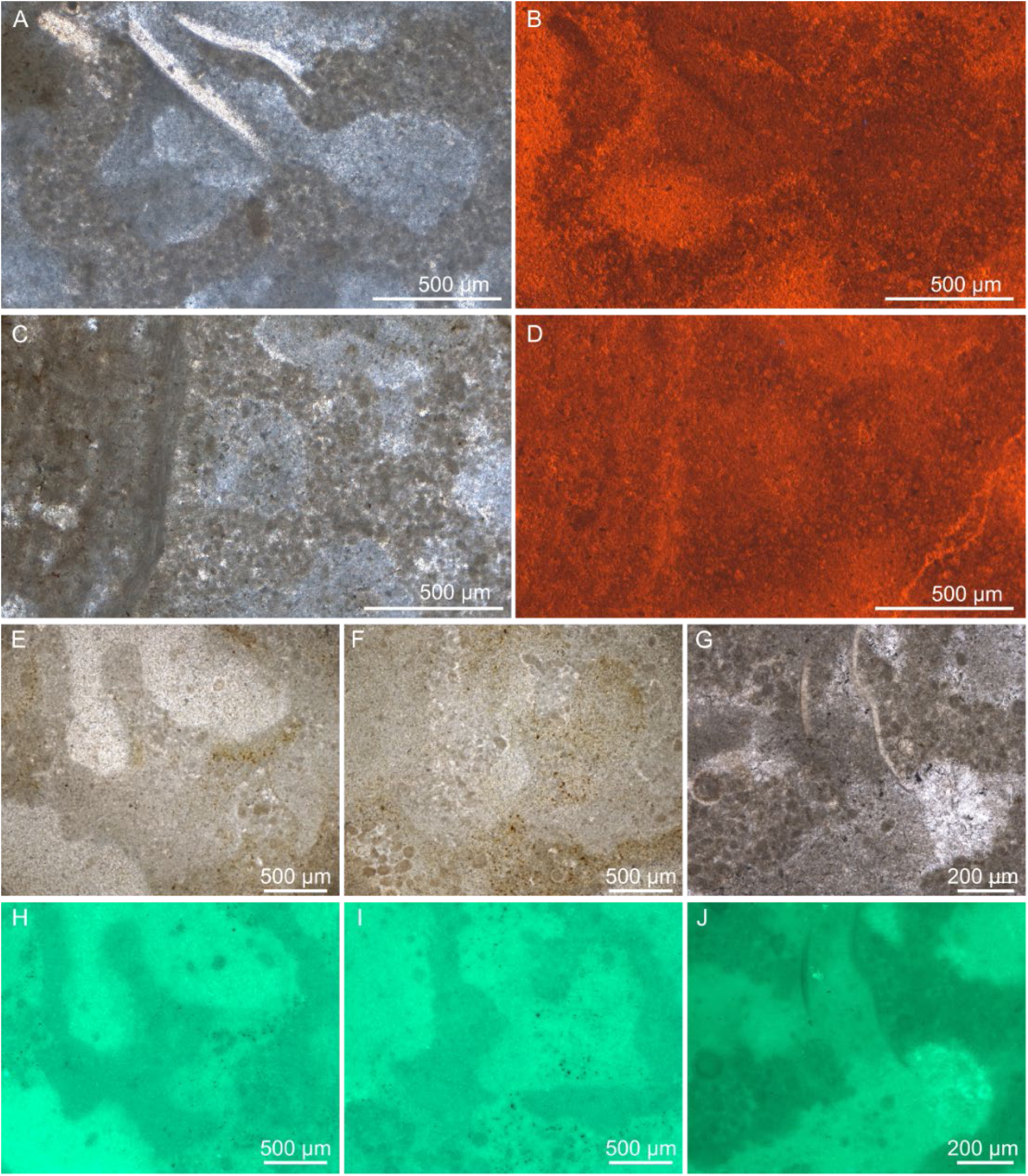
Cathodoluminescence (CL) and fluorescence petrography of the studied materials. (A–D) Corresponding areas containing the labyrinthine partitions in plane polarized light (PPL)(A, C) and under a CL microscope (B, D). (E–J) Corresponding areas containing the labyrinthine partitions in PPL (E, F, G) and under a fluorescence microscope with blue incident light (H, I, J). Thin section numbers: xz-05a-4 for A–D; xz-02 for E, F, H, I; xz-05b-1 for G and J.

The slab analyzed using serial grinding and 3-D reconstruction is a part of SPF-1, and it reveals that the labyrinthine partitions are three-dimensionally anastomosing. The anastomosed elements are typically platy, with only local occurrences of filamentous elements (Fig. 7B, C, E–I). In the 3-D model, the analyzed fossil is in a bowl shape and widens upwards (Fig. 7E). Three cavities with diameters up to 2 mm are recognized (Fig. 7A, D–I). The morphology and the nature of these cavities will be discussed in detail in Section 4.1.3.

**Fig. 7.**
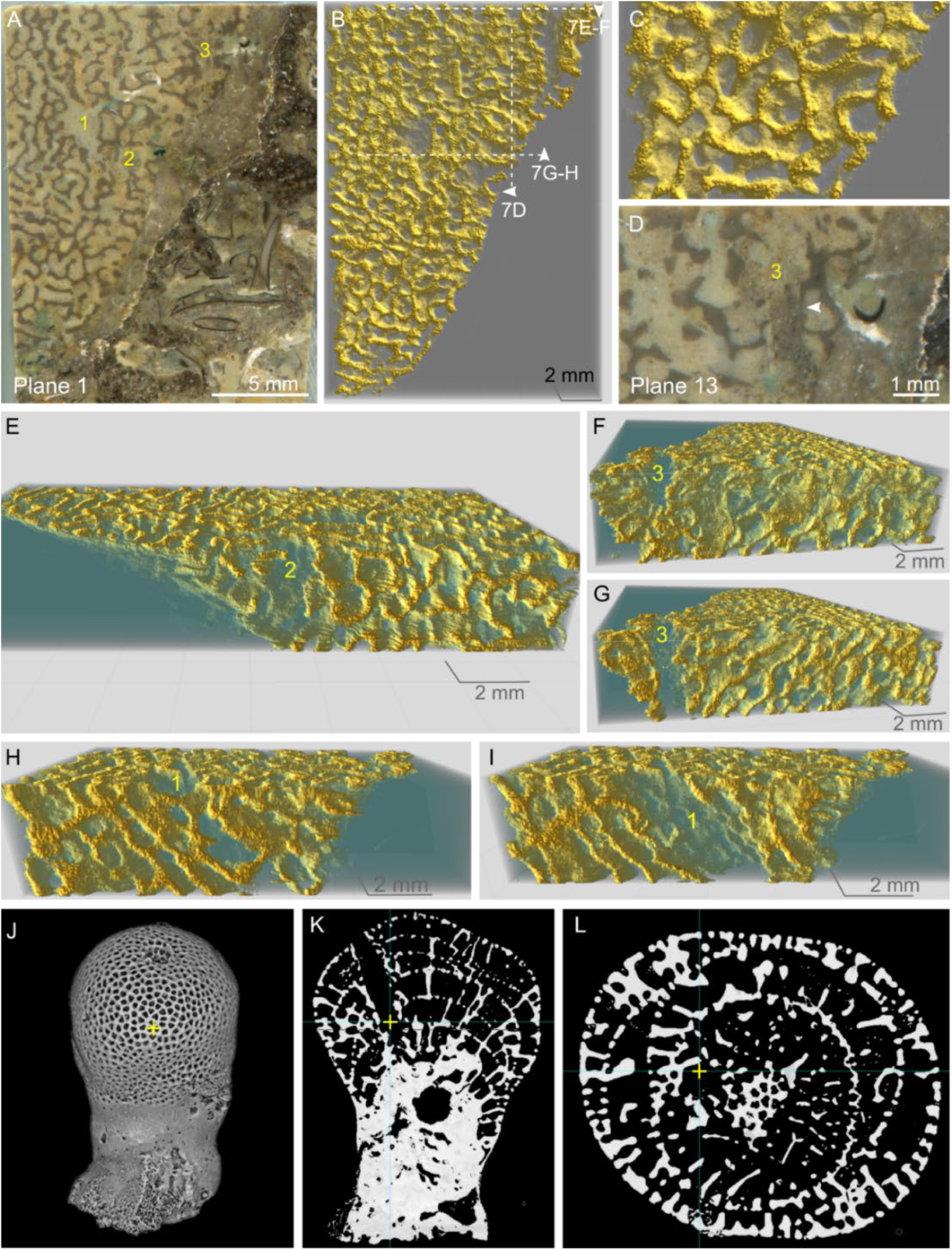
Three-dimensional reconstruction of a part of SPF-1. (A) The topmost plane of the serial grinding, numbers indicating the three cavities discussed in the main text. (B) The rendered 3-D structure of the analyzed volume, in the same orientation as A. Dashed lines and arrows indicate the position and viewing direction of the images in Fig. 7E–I, respectively. (C) A close view of B, showing the anastomosing platy partitions. (D) Cavity-3 in plane 13. The white arrow indicates possible constrained growth of the partition. (E) A cross-section of the 3-D model showing the morphology of cavity-2. (F and G) Cross-sections of the 3-D model showing the morphology of cavity-3. G is further clipped than F. (H and I) Cross-sections of the 3-D model showing the morphology of cavity-1. I is further clipped than H. (J–L) Three-dimensional structure of a *Vaceletia* specimen (J) and its two cross-sections (K and L). The yellow cross marks the same position in different views. The full length of the specimen in J is 7 mm.

## 4. Discussion

### 4.1. The sponge affinity of the labyrinthine fossil

In hand specimens and thin sections, the labyrinthine fossils are reminiscent of some coralline/hypercalcified sponge (e.g., stromatoporoids of the order Stromatoporida) (Stearn, 2015a), the enigmatic terminal Ediacaran fossil *Namapoikia* (Mehra et al., 2020; Wood and Penny, 2018; Wood et al., 2002), and the maceriate microbialite *Favosamaceria* (Shapiro and Awramik, 2006). All these structures and fossils exhibit a labyrinth pattern in plan view and elongated, branching and jointing elements in vertical cross-section.

#### 4.1.1. Comparisons by general morphology

In the first place, the microbialite *Favosamaceria* can be differentiated from other fossils because of its huge dimension, 1–2 orders of magnitude larger than the others (Table 1). Indeed, *Favosamaceria* is defined by its architecture presented at the decimetric level (macrostructure) (Shapiro and Awramik, 2006). However, the maceria of *Favosamaceria* is composed of packed peloids, and the interspaces are filled by micrite and sparite. This composition is similar to that of the Xuzhou fossil. *Favosamaceria* is a microbialite that is believed to have been formed under significant biological control, but the producers and processes remain unknown (Shapiro and Awramik, 2006). *Favosamaceria*-like microbialite macrostructures were also found being constructed by microstromatolites, calcimicrobes, and possible non-spicular sponges (Lee and Riding, 2022).

**Table 1.** Morphological comparisons among *Favosamaceria*, *Namapoikia*, two stromatoporoid species, and the Xuzhou fossil.

|  | *Favosamaceria* | *Namapoikia* | *Stromatopora*<br>*hüpschii* | *Syringostromella*<br>*cf. labryrinthea* | Xuzhou<br>fossil |
| --- | --- | --- | --- | --- | --- |
| Thickness of<br>partitions/maceria/skeletons<br>(mm)* | 10–100 | 0.20–1.65 | 0.1–0.2 | 0.1–0.2 | 0.1–0.23 |
| Interspace width (mm) | 10 | 0.20–2.91 | 0.1–0.2 | 0.1–0.2 | 0.3–0.6 |
| Distance between the<br>branching points of<br>partitions/maceria/skeletons<br>in vertical cross-sections<br>(mm) | 20–300 | NA | NA | NA | 0.4–0.9 |
| Aquiferous canals | Absent | Absent | NA | NA | Present |
| Horizontal partitions | Absent | Often absent | Present<br>(dissepiments) | Present<br>(dissepiments) | Absent |
| Distinct boundary against<br>surrounding microbialites | NA | Present | NA | NA | Present |
| References | Shapiro and<br>Awramik (2006) | Mehra et al.<br>(2020); Wood<br>and Penny<br>(2018) | Huang et al.<br>(2024) | Huang et al.<br>(2024) | This study |

\* The labyrinthine walls in *Favosamaceria* were termed “maceria”(Shapiro and Awramik, 2006), and those in *Namapoikia* were called “partition” or “skeleton” by different authors (Mehra et al., 2020; Wood and Penny, 2018).

The affinity of *Namapoikia* is controversial. Wood and Penny (2018) described transverse tabulae bridging the vertical walls and highlighted the similarity of this skeletal architecture to that of stromatoporoid and chaetetid-grade sponges. However, tabulae were not observed in the specimens studied by Mehra et al. (2020). Instead, the latter authors negated the poriferan affinity of *Namapoikia* because of its thick walls, wide interspaces, and the large variance of partition and interspace sizes, all unseen in other Phanerozoic sponges (Mehra et al., 2020). Nevertheless, a microbialite affinity of *Namapoikia* has difficulties in interpreting the consistent skeletal composition of this fossil and its distinct boundary against the surrounding microbial fabrics (Wood and Penny, 2018; Wood et al., 2002).

Like *Namapoikia*, the studied labyrinthine fossils from Xuzhou are closely associated with microbialites but always maintain a clear boundary against the latter. This is similar to the growth habit of sponges dwelling in microbialites from other Phanerozoic localities, e.g., *Gravestockia pharetroniensis* from the lower Cambrian of Australia (Reitner et al., 2017) and the suggested non-spicular sponge fossils from Triassic and Carboniferous stromatolites (Luo and Reitner, 2016, 2014; Pei et al., 2021). Meanwhile, the dimension of the labyrinthine fossils falls into the range of common hypercalcified sponges (Table 1; Mehra et al., 2020). In regard to the partition architecture, these fossils are most similar to some stromatoporoid taxa such as *Stromatopora* and *Syringostromella*, whose skeletons are dominantly built with pachysteles and pachystromes (Huang et al., 2024; Stearn, 2015a, 2015b). Nevertheless, the Cambrian labyrinthine fossils do not have dissepiments and show wider interpartition spaces.

#### 4.1.2. Comparisons by composition

Another significant difference between the labyrinthine fossils and stromatoporoids is their composition. Stromatoporoid skeleton is typically composed of biomineralized carbonates with consistent microstructures (Stearn, 2015c). In contrast, the partitions of the labyrinthine fossil dominantly consist of packed detritus particles, supplemented by micrites in some places. However, if considering the large variety of methods that sponges can use to build their skeletons, this question of compositional difference may become understandable.

The massive calcareous basal skeleton of coralline/hypercalcified sponges can be formed either by progressively cementing and enveloping primary spicules or by biologically controlled mineral precipitation unrelated to spicules (Vacelet et al., 2015). The second method, in turn, includes multiple ways of biomineralization. *Astrosclera willeyana* initially secretes sclerodermites intracellularly, then transports them to the growing basal skeletons and leaves them to grow outside of the cell (Gautret, 1986; Wörheide 1997). *Acanthochaetetes wellsi* accretes the microlamillae of its basal skeleton by inducing carbonate precipitation within a thin zone (300–500 nm) below its basopinacoderm (Reitner and Gautret, 1996). The carbonate skeleton of *Vaceletia* was formed because of direct aragonite precipitation in the organic skeletal fibers (Reitner, 1992; Reitner et al., 1997; Vacelet, 1979). This last type of biomineralization results in skeletons with microgranular structures and was regarded as a model to interpret the biomineralization of archaeocyaths (Reitner et al., 1997). This biomineralization mode is also applicable to interpret the micritic part of the partitions in the labyrinthine fossils.

*Vaceletia* is a non-spicular demosponge belonging to the order Dictyoceratida (Erpenbeck et al., 2020; Wörheide, 2008). Although *Vaceletia* does not, many taxa in this order are known to incorporate and agglutinate foreign particles. For instance, the skeleton of *Dysidea arenaria* (order Dictyoceratida) is “made from sand and abundant spicule fragments, bound by scarce spongin” (p. 600, Pulitzer-Finali and Pronzato, 1999). Indeed, beyond the order Dictyoceratida, incorporating allochthonous grains into the skeletons is not rare throughout the whole demosponge class (Schönberg, 2016). For example, many genera in the family Chondropsidae (order Poecilosclerida) partly or wholly replace the spicule bundles with agglutinated detritus (van Soest, 2002). It is noteworthy that *Psammoclema*, although belonging to a spicular order and indeed secretes siliceous microscleres, exhibits a skeleton composed of agglutinated particles only (van Soest, 2002; Wiedenmayer, 1989).

With these facts in mind, it is conceivable that the labyrinthine fossils may have been an organism similar to *Dysidea arenaria* or *Psammoclema*, living with the habit of constructing agglutinated skeletons. Meanwhile, the organic skeleton of this fossil organism is similar to *Vaceletia* in being able to be biomineralized with micrites. The proportion of detritus versus biominerals in the skeleton may have been controlled by the environmental detritus supply and/or by the organismal regulation. When the detritus supply is low, the organism may produce a spongin-dominant skeleton, which is then mineralized by micrites. In contrast, when the sedimentation rate is high, the sponge could also grow faster by building new skeletons with packed foreign detritus. It is notable that this is not the first time we see sediment incorporation in a coralline/hypercalcified sponge. The Jurassic stromatoporoid *Calciagglutispongia yabei* Reitner, 1992 was found incorporating detritus into the center of its calcified skeletal fibers.

Finally, although the organic skeletons of most extant demosponges are fibrous, there are examples whose skeletons are anastomosing and platy (*Ianthella basta*, Fig. 11 in Pulitzer-Finali and Pronzato, 1999).

#### 4.1.3. Aquiferous canals

As mentioned in Section 3.3, the 3-D reconstruction revealed three cavities in the studied specimen. Among them, cavity-1 is located in the center of the analyzed slab and is best preserved among the three (Fig. 7A). This cavity vertically penetrates the analyzed specimen and is connected with inter-partition spaces (Fig. 7A, B, H, I; Supplementary material 1). The partitions around the cavity do not show evidence of destruction, while the shape of the cavity cross-section rapidly changes within the analyzed vertical distance of 3 mm (for comparison, the cavity diameter is around 2 mm) (Supplementary material 1). These features exclude a burrow interpretation of this cavity. Instead, the orientation of this cavity, as well as the partitions and interspaces around it, is analogous to what is observed around the siphon (spongocoel) of *Vaceletia* (Fig. 7J–L).

Cavity-2 is similar to cavity-1 in diameter, orientation, and the lack of evidence of destruction (Fig. 7A, B, E; Supplementary material 1). However, it is located on the edge of the analyzed slab, and only a small corner of it is observable in the analyzed data. Therefore, we cannot determine its nature as confidently as for cavity-1.

Cavity-3, at first glance, looks like a mechanically induced fissure in the fossil (Fig. 7A and B). Indeed, detrital particles are more densely accumulated in this cavity than in other parts of the fossil (Supplementary material 1B), and the serial grinding planes 12, 13, 14 do indicate an inhibited growth or a destruction of the partition next to the detrital patch (Fig. 7D; Supplementary material 1A and B). However, one end of this “fissure” is closed by undisturbed partitions (Fig. 7B, F). It’s possible that cavity-3 represents a mechanically altered or passively filled canal in the early growth stage of the organism.

### 4.2 Systematic Paleontology

Taking these discussions in section 4.1 together, a sponge affinity seems to be the most reasonable interpretation for all the characteristics of the labyrinthine fossils, including morphology, partition composition, inner cavities, and interaction with the ambient microbial systems. For the convenience of future investigation and discussion, here we provide a formal taxonomic description for this new fossil, and in the following text, the partitions will be called “skeletons”.

Phylum Porifera Grant, 1836

Class Demospongiae Sollas, 1885

Order Uncertain Family Uncertain

Genus *Psammolectospongia* gen. nov.

Etymology – “psammo-” is a Latin prefix meaning sand; “lecto-” means collecting; “spongia” means sponge.

Type species – *Psammolectospongia beiwangensis* gen. et sp. nov.

Diagnosis – An encrusting sponge with a skeleton dominantly composed of tightly packed, agglutinated, silt-to sand-sized detritus. A minor part of the skeleton may be composed of micrites. The skeletal frame consists of anastomosed platy elements, which are mainly oriented subparallel to the top-down axis of the organism. The platy elements are around 0.1–0.2 μm thick, with interspaces of about 0.3–0.6 μm wide. Multiple exhalant aquiferous canals of 1–2 mm wide can be present in each individual.

Remarks – This fossil is tentatively assigned to the class Demospongiae because, among known extant taxa, demosponges are the only group that has been observed to incorporate detritus into the body (Schönberg, 2016). However, the order-and family-level taxonomy is difficult to determine, since detritus incorporation has been observed in multiple spicular and non-spicular demosponge orders (Section 4.1.2).

*Psammolectospongia beiwangensis* gen. et sp. nov. (Fig. 1F–J, 2A, 3A, 4A–H, 5A–I, 6A–G, 7A–I)

Etymology — “beiwang*-*” is the name of the village where the described fossils were found. Diagnosis — As the diagnosis of the genus.

Description — As in Section 3.3.

Materials — Six specimens processed into thin sections and slabs. Holotype, SPF-1, including 2 thin sections, 2 slabs, and 1 serial grinding dataset. Paratypes: SPF-2 (6 thin sections), SPF-3 (1 thin section), SPF-4 (1 thin section and 1 slab), SPF-5 (1 slab), SPF-6 (1 slab).

Occurrence — The new species is so far only found in the *Redlichia nobilis* Zone, Cambrian Stage 4, in the Beiwang Village, Xuzhou City, China.

### 4.3 Preservation of the agglutinated sponge skeletons

Cathodoluminescence (CL) microscopy and fluorescence microscopy have both been proven helpful tools in the study of carbonate diagenesis since the second half of the 20^th^ century (Dravis and Yurewicz, 1985). As described in Section 3.3, neither method revealed significant diagenetic alteration in *Psammolectospongia beiwangensis* gen. et sp. nov. and its host fabrics.

Compared with CL, fluorescence microscopy appears to be more sensitive in detecting the boundary between the *P. beiwangensis* skeleton and the interspace fillings, even when the two parts look indistinguishable in transmitted light. Regardless of a micritic or detrital composition, the *P. beiwangensis* skeleton always appears to be darker than the interspace fillings. However, this does not mean that the skeleton was abiotic. As a comparison, skeletal fragments (including trilobites, echinoderms, and brachiopods) and *Girvanella* fossils in the studied materials can be variously dull or bright (e.g., dull ones shown in Fig. 6J). The Holocene corallites described in Dravis and Yurewicz (1985) are also non-fluorescent. Indeed, the fluorescence intensity of geological materials is not solely determined by the amounts and types of remaining organic matter, but also affected by the relative concentration of activators and quenchers, e.g., Mg activates blueish-green fluorescence, while Fe quenches fluorescence (Rost, 1995).

Regardless of the cause of the fluorescence pattern, the different fluorescent behaviors between the *P. beiwangensis* skeleton and the interspace fillings are stable. Part of the sparitic cements between agglutinated detritus are fluorescent, but there are also non-fluorescent cements in the skeletons (Fig. 6E–J). It is possible that the non-fluorescent sparitic cements were transformed from micrites by neomorphism. That is to say, the micritic biomineralization in *P. beiwangensis* may have also played a role in holding the agglutinated detritus together.

## 5. Conclusion

After comparing with fossil and extant sponges, as well as similar microbialites and enigmatic fossils, the labyrinthine fossils found in the microbialites of Cambrian Stage 4 of Xuzhou seem to fit the affinity of an aggutinating demosponge best. This is the first confirmed example of early Paleozoic sponges that are capable of actively incorporating sediment into the primary skeletons. This finding indicates that a high physiological flexibility, even in regard to skeletal composition, has been developed in shallow water sponges since the early Cambrian.

**Supplementary Material 1.** Raw data and video of the 3-D reconstruction (all available at https://doi.org/10.57760/sciencedb.28369). (A) the original scans of the serial grinding; (B) the stack of aligned images; (C) the segmented stack with background removed; (D) a modified version of the segmented stack, with interpolations to adjust its thickness according to the measured real dimension; (E) the Voreen workplace to render the stack in D and to generate the video; (F) a video of the 3-D image.

## Acknowledgements

We thank Wen-Qian Wang (Nanjing University) and Jing Feng (NIGPAS) for allowing access to the CL and the fluorescent microscopy, respectively. Zhi-Kang Kou (NIGPAS) and the deceased You-Dong Chen (NIGPAS) prepared the thin sections, and Le Yang (NIGPAS) and Ruo-Dan Chen (freelancer) helped with the acquisition and processing of the serial grinding data. This study was supported by the National Natural Science Foundation of China (Grant No. 42330209).

